# Cognitive Functions Mediate the Effect of Preterm Birth on Mathematics Skills in Young Children

**DOI:** 10.1101/868547

**Authors:** Julia Anna Adrian, Roger Bakeman, Natacha Akshoomoff, Frank Haist

**Affiliations:** Department of Cognitive Science, UC San Diego; Center for Human Development, UC San Diego; Department of Psychology, Georgia State University; Department of Psychiatry, UC San Diego

**Keywords:** Preterm birth, mathematics, cognitive functions, phonological processing, mediation, development

## Abstract

Children born preterm are at risk for cognitive deficits and lower academic achievement. Notably, mathematics achievement is generally most affected. Here, we investigated the cognitive functions mediating early mathematics skills and how these are impacted by preterm birth. Healthy children born preterm (gestational age at birth < 33 weeks; *n* = 51) and children born full term (*n* = 27) were tested at ages 5, 6, and 7 years with a comprehensive battery of tests. We categorized items of the TEMA-3: Test for Early Mathematics Abilities Third Edition into number skills and arithmetic skills. Using multiple mediation models, we assessed how the effect of preterm birth on mathematics skills is mediated spatial working memory, inhibitory control, visual-motor integration, and phonological processing. Both number and arithmetic skills showed group differences, but with different developmental trajectories. The initial poorer performance observed in the preterm children decreased over time for number skills but increased for arithmetic skills. Phonological processing, visual-motor integration, and inhibitory control were poorer in children born preterm. These cognitive functions, particularly phonological processing, had a mediating effect on both types of mathematics skills. These findings help define and chart the trajectory of the specific cognitive skills directly influencing math deficit phenotypes in children born very preterm. This knowledge provides guidance for targeted evaluation and treatment implementation.

---

Preterm birth (before 37 weeks of gestation) occurs in about 10% of all live births (Chawanpaiboon et al., 2019) and can be associated with brain injury and other health issues (Ramachandrappa & Jain, 2009; Volpe, 2009). Advances in neonatal medical care have improved survival rates and severity of health outcomes (Grytten et al., 2017). Nevertheless, even in the absence of severe disabilities, children born preterm often suffer from developmental, cognitive, and behavioral problems (Anderson, 2014).

Children born preterm before 33 weeks of gestation are especially at increased risk for deficits in cognitive functions and academic achievement (Johnson, Wolke, Hennessy, & Marlow, 2011), specifically in mathematics (Aarnoudse-Moens, Oosterlaan, Duivenvoorden, van Goudoever, & Weisglas-Kuperus, 2011; Akshoomoff et al., 2017; Taylor, Espy, & Anderson, 2009). Lower educational outcomes following this level of preterm birth have been reported in early school age (Pritchard et al., 2009; Taylor et al., 2018), adolescence (Litt et al., 2012; Rose, Feldman, & Jankowski, 2011), and adulthood (Løhaugen et al., 2010). Little is known about the developmental trajectory of emerging mathematics skills in preterm children.

In the general population, school-entry mathematics skills are a strong predictor of later academic achievement (Duncan et al., 2007). Middle-school mathematics skills have been shown to mediate the relationship between preterm birth and adult wealth (Basten, Jaekel, Johnson, Gilmore, & Wolke, 2015). Here we focus on the development of mathematics skills in the critical early school-age period.

## Mathematics Comprises Multiple Skills

One major issue regarding the study of children’s early mathematics development is that it is often seen as one skill, rather than the variety of different skills that constitute mathematics ability. Standardized achievement tests used with children and adults are typically designed to span a wide age range, sample a broad array of academic skills based on age expectations, and provide one overall score (e.g. Woodcock, McGrew, & Mather, 2007). In younger children, the Test for Early Mathematics Ability, Third Edition (TEMA-3, Ginsburg & Baroody, 2003) has been used in studies of typically and atypically developing children (Fuhs & McNeil, 2013; Hasler & Akshoomoff, 2019; Kull & Coley, 2015; Mazzocco, Feigenson, & Halberda, 2011; Schneider et al., 2017). While most studies used the overall scaled *Mathematics Ability Score* to assess children’s performance, the TEMA-3 also provides raw scores of “informal” and “formal” mathematics abilities. Informal mathematics abilities include numbering, counting, magnitude comparisons, and using fingers or other markers to solve simple arithmetic problems. In comparison, formal math skills are abilities learned in school including the understanding and use of numerals, exact magnitude specification, and memorized facts for addition, subtraction, and multiplication. Libertus, Feigenson, & Halberda (2013) found that young children’s numerical approximation abilities predicted their informal but not formal mathematics abilities. While the TEMA-3 informal and formal distinction is useful, it is based on descriptive face-valid qualities. In this study, we followed the guidance from Ryoo and colleagues (2015) to create categories of specifically defined math skills. Two types of mathematics skills were defined (see Table 1): (a) *number skills*, including items that are related to the ordering based on numerical magnitudes such as counting and number comparison skills (28 items), and (b) *arithmetic skills*, including items that are related to manipulation of numbers such as calculation skills with problems that are presented in story form or via equations (36 items). While test items in these categories require distinct sets of skills, number and arithmetic skills do not develop independent from each other. During early school age, arithmetic skills have been shown to be predicted by (particularly symbolic) number skills (Lyons, Price, Vaessen, Blomert, & Ansari, 2014).

**Table 1.** Definition of Number Skills and Arithmetic Skills.

|  | Category† | # items | Type of task |
| --- | --- | --- | --- |
| **Number Skills** | Verbal Counting | 14 | Counting until a certain number; tell the successor number |
|  | Counting Objects | 7 | Counting objects |
|  | Numerical Comparison | 7 | Which number is more; which number is closer to a third number |
| **Arithmetic Skills** | Set Construction | 9 | Division problems presented in story/money form |
|  | Calculation | 18 | Calculation problems presented in story form or abstract |
|  | Number Facts | 9 | Single digits addition (verbal, no story form, with time limit) |
| Not included | Numeral Literacy | 8 | Reading & writing numerals |

*Note.* Number skills are measured by TEMA-3 test items that require ordering of numbers by magnitude. Arithmetic skills are measured through TEMA-3 test items that require manipulation of numbers.
†Category as defined by Ryoo et al. (2015)

## Distinct Cognitive Functions Contribute to Mathematics Skills

Mathematics performance requires the integration of a complex set of skills. Previous work has identified working memory and inhibitory control, visual-motor integration, and phonological processing as critical components contributing to mathematics abilities (De Smedt et al., 2010; Geary & Moore, 2016; Kulp, 1999). A specific cognitive function might be more important for one type of mathematics skill than for another one. For example, Lan, Legare, Ponitz, Li, & Morrison (2011) found that working memory uniquely predicted calculation skills in preschoolers, while counting skills were predicted both by working memory and inhibition. Dividing mathematics skills into number and arithmetic skills thus provides the opportunity to study the influence of cognitive functions on specific mathematics skills, particularly those functions that are impacted by preterm birth.

Executive functions are a robust predictor of mathematics skills in full-term and preterm children. In a longitudinal study of over 1200 typically developing children, Ribner, Willoughby, & Blair (2017) found that executive function skills at age 5 strongly predicted mathematics achievement in 5^th^ grade. Working memory might underpin mathematics skills when it is necessary to mentally retain and retrieve relevant information. Inhibitory control may contribute more to the suppression of inappropriate but mentally prevalent answers or strategies (Bull & Lee, 2014). Rose et al. (2011) suggested a *cascade of effects* from prematurity to slower processing speed, to poorer executive functions (working memory), and finally lower academic achievement, after examining 11-year-old preterm children with a birth weight of< 1750g. Executive function deficits were also reported in 3–5 year old preterm children and found to meditate the effect of gestational age on behavioral problems (Loe, Feldman, & Huffman, 2014).

Visual-motor integration has not been studied extensively but is a potential mediator between preterm birth and low mathematics performance because preterm children have an increased risk of visual-motor integration deficits (Geldof, van Wassenaer, de Kieviet, Kok, & Oosterlaan, 2012).. Deficits in visual-motor integration have been shown to be associated with lower mathematics performance in typically developing children (Sortor & Kulp, 2003). A recent study of children born extremely preterm (before 28 weeks gestation) showed involvement of visual-motor integration in mathematics performance (Taylor et al., 2018).

In addition, verbal skills contribute to mathematics performance. Language functions are involved when solving mathematics problems, for example through representation and manipulation of magnitudes in form of number words. Specifically, the extant literature shows that phonological processing makes a specific contribution to mathematics abilities. Phonological processing includes the ability to perceive, produce, discriminate, and manipulate specific sounds of a language. In a typically developing cohort, phonological difficulty at age 5 was associated with deficits in formal mathematics components at age 7 (Jordan, Wylie, & Mulhern, 2010). Phonological awareness is a specific part of phonological processing that describes the ability to concatenate and remove phonological segments to form words. De Smedt, Taylor, Archibald, & Ansari (2010) reported a specific and unique association between phonological awareness and single-digit arithmetic skills in typically developing children at age 10. Deficits in phonological awareness, phonological processing, and other language outcomes have been reported in preterm children (Vohr, 2014), though fewer studies have investigated the link between phonology and mathematics in preterm children.

Most studies investigating mathematics performance in full-term and preterm children examine overall performance on standardized mathematics tests, which include a variety of different skills. It is thus difficult to understand which specific mathematics skills are affected at different ages. It remains an open question whether number and arithmetic skills are differentially affected by preterm birth and if other cognitive functions, such as working memory, inhibitory control, visual-motor integration, and phonological processing are related to these mathematics skills. We chose to examine these particular cognitive functions based on recent research linking them to mathematics skills in children who were born preterm (Hasler & Akshoomoff, 2019; Tatsuoka et al., 2016; Taylor et al., 2018; van Veen, van Wassenaer-Leemhuis, van Kaam, Oosterlaan, & Aarnoudse-Moens, 2019). Furthermore, studies that investigate the effect of cognitive functions on mathematics skills often only study one of those functions (e.g. working memory) at a time. This does not allow us to understand how large the effects of these cognitive functions are in comparison to one another.

Our study first compared the status of number skills and arithmetic skills in children born preterm and full-term. Next, using mediation analysis, we determined which cognitive functions mediated the relationship between preterm birth and mathematics skills. Our study addresses limitations in prior research by including multiple cognitive functions in the same model, namely working memory, inhibitory control, visual-motor integration, and phonological processing.

## Method

### Participants

We recruited 51 children born preterm (24–32 weeks gestational age) and 27 children born full-term (38–41 weeks gestational age). Participant characteristics are summarized in Table 2. By definition, the preterm group had a significantly lower gestational age at birth and birth weight. There were no significant group differences in terms of sex, age at testing, household income, and race. Maternal education was lower in children born preterm compared to full-term.

**Table 2.** Participant characteristics.

|  | Preterm<br>n = 51 | Full-term<br>n = 27 | Effect size<br>(Test statistic, p-value) |
| --- | --- | --- | --- |
| GA at birth in weeks<br>mean (SD, min-max) | 29.4<br>(2.00, 25-32) | 39.7<br>(0.77, 38-41) | $\eta^2=.894$<br>(F=643, $p<.001$) |
| Birth weight in g<br>mean (SD, min-max) | 1327<br>(439, 680-2410) | 3410<br>(561, 2353-4422) | $\eta^2=.806$<br>(F=316, $p<.001$) |
| Sex (female) | 47% | 48% | $V=.010$ ($\chi^2=.008$, $p=.927$) |
| Age at testing: mean (SD) |  |  |  |
| Time 1 (Age 5) | 5.3 (0.3) | 5.4 (0.3) | $\eta^2=.018$ (F=1.37, $p=.245$) |
| Time 2 (Age 6) | 6.4 (0.4) | 6.4 (0.3) | $\eta^2=.006$ (F=.421, $p=.518$) |
| Time 3 (Age 7) | 7.3 (0.4) | 7.4 (0.3) | $\eta^2=.005$ (F=.420, $p=.519$) |
| Maternal education<br>absolute (relative frequency) |  |  |  |
| 1) high school degree | 7 - 14% | 3 - 10% | $\eta^2=.048$ |
| 2) 1–3 year college | 18 - 35% | 6 - 21% | (U=514, $p=.055$) |
| 3) college degree | 21 - 41% | 10 - 35% |  |
| 4) graduate degree | 5 - 10% | 10 - 35% |  |
| Household income:<br>absolute (relative frequency) |  |  |  |
| 1) below \$50,000 | 8 - 16% | 1 - 3% | $\eta^2=.002$ |
| 2) \$50,000 – <\$100,000 | 16 - 31% | 13 - 45% | (U=621, $p=.736$) |
| 3) \$100,000 – <\$200,000 | 18 - 35% | 13 - 45% | |
| 4) \$200,000 and above | 8 - 16% | 1 - 3% | |
| Missing information | 1 - 2% | 1 - 3% |  |
| Race<br>absolute (relative frequency) |  |  |  |
| African American | 1 - 2% | 0 - 0% | $V=.281$ |
| Asian | 2 - 4% | 4 - 15% | ($\chi^2=6.15$, $p=.188$) |
| Caucasian | 40 - 78% | 16 - 59% |  |
| Mixed/other | 7 - 14% | 7 - 26% |  |
| Missing information | 1 - 2% | 0 - 0% |  |
| Ethnicity (Hispanic/Latinx) | 19 - 37% | 8 - 30% | $V=.28$ ($\chi^2=.454$, $p=.501$) |

*Note.* Group comparisons were performed via ANOVA F[1,77], effect size: partial $\eta^2$ for GA at birth, birth weight, and age at testing; via Mann-Whitney U test, effect size: $\eta^2$ for maternal education, and household income; and via Chi-square test, effect size: Cramer's V for sex, race, and ethnicity. GA: gestational age; g: grams..

The preterm participants were recruited primarily from the UC San Diego High-Risk Infant Follow Up Clinic. Inclusion criteria for the preterm sample were gestational age at birth of < 33 weeks and absence of severe congenital, physical or neurological disabilities. Children were excluded if they had a history of severe brain injury (intraventricular hemorrhage of grade 3-4, cystic periventricular leukomalacia, moderate-severe ventricular dilation), genetic/chromosomal abnormalities affecting development, severe disability (e.g. bilateral blindness, cerebral palsy), or acquired neurological disorders unrelated to preterm birth. Of the 51 children born preterm, 10 were born extremely preterm (< 28 weeks of gestation); additionally, 16 children were born with very low birth weight (1000g–1500g) and 17 with extremely low birth weight (< 1000g). Six preterm participants had intraventricular hemorrhage of grade 1-2 (later resolved), none had periventricular leukomalacia, five had bronchopulmonary dysplasia, and five were small for gestational age.

The full-term participants were recruited through the Center for Human Development at UC San Diego. Inclusion criteria for the full-term sample were gestational age at birth of > 37 weeks and no history of neurological, psychiatric, or developmental disorders. Additionally, all participants were required to be native English speakers. As the data of this study is part of a larger project that includes MRI imaging, participants were excluded if they had a history of anxiety and/or metal implants that would interfere with scanning. The Institutional Review Board at UC San Diego approved the study. Legal guardians gave written informed consent, and children of age 7 years and older gave assent.

### Design and Procedure

We used a longitudinal design with each child receiving a comprehensive battery of cognitive and academic tests, health and demographic questionnaires and MRI imaging at three time points at approximate ages of 5, 6, and 7 years. Baseline testing was performed within six months of starting kindergarten, age at testing across the three assessments is described in Table 2. Partial behavioral and MRI results are described elsewhere (Hasler & Akshoomoff, 2019; Hasler, Brown, & Akshoomoff, 2019).

### Assessment of Mathematics Skills

We assessed mathematics skills using the Test for Early Mathematics Ability, 3^rd^ Edition (TEMA-3; Ginsburg & Baroody, 2003). The TEMA-3 is designed for children between 3 and 8 years old and includes72 items that a subset may be given based on age and performance. Overall performance is expressed in sum raw score of correct items and the standardized Mathematics Ability Score (mean=100, SD=15). These measures are commonly used to characterize ‘mathematics ability’ in children. Ryoo et al. (2015) grouped the 72 test items from the TEMA-3 into seven categories: Verbal Counting, Counting Objects, Numerical Comparison, Set Construction, Calculation, Number Facts, Numeral Literacy that were validated by confirmatory factor analysis from their sample of 389 children. Their factor structure fit the longitudinal data better than the “formal” and “informal” TEMA-3 mathematics dichotomy. While the Ryoo et al. (2015) categorization scheme may be more theoretically useful, each of the seven categories contained only a small number of items and may be vulnerable to statistical instability. Furthermore, some of the items in different categories require the same type of skill. For example, items from the categories Set Construction, Calculation, and Number Facts all require the manipulation of numbers (arithmetic). Similarly, items in the Verbal Counting, Counting Objects, and Numerical Comparison clusters require ordering numbers based on the magnitudes they represent.

Based on the types of skills required to solve the problems, we clustered the TEMA-3 items by combining categories from Ryoo et al. (2015) into *number skills* and *arithmetic skills* (see Table 1). The category number skills comprises 28 items from the subcategories Verbal Counting, Counting Objects, and Numerical Comparison as defined by Ryoo et al. (2015). These items require the participant to order quantities by magnitude. The arithmetic skills category includes the 36 items from the subcategories Set Construction, Calculation, and Number Facts. These items require the participant to manipulate numbers to solve abstract and concrete (story-form) problems. One subcategory, Numeral Literacy, required participants to read and write numbers, a skill different from the other subcategories. These items were excluded from the present analyses.

### Assessment of Cognitive Functions

#### Spatial working memory

We assessed spatial working memory via the Cambridge Neuropsychological Testing Automated Battery (*CANTAB® Cognitive assessment software*, 2017) Spatial Working Memory Task (SWM). This task is designed for participants from 4 to 99 years of age. It is a nonverbal, computerized task presented on a touch screen. The screen shows colored squares, the participant has to find a token that is hidden behind one of them. Once the token is found it is hidden again behind one of the squares under which it was not previously hidden. The participant has to use spatial working memory to successfully and efficiently find the hidden token and not search under the same square twice. Task difficulty increased by increasing the number of squares. Task performance is measured inversely through the number of errors made (number of squares that are searched multiple times, “between errors”). To simplify interpretation of the results, the inverse of this error measure is used for analysis, such that a more positive measure of SWM corresponds to better spatial working memory.

#### Inhibitory control

We assessed inhibitory control via the CANTAB Stop Signal Reaction Task (SST), designed for ages 4 to 99 years. It is a ‘go/no-go’ style nonverbal, computerized task presented on a touch screen. The participants see a circle in the center of the screen and one rectangle on either side of it. When an arrow appears on the screen, the participant’s task is to touch the rectangle to which the arrow points as fast as possible. If they hear the auditory stop signal the participant has to inhibit their response and not touch the screen. Task performance is measured through the stop signal reaction time (SSRT). To simplify interpretation of the results and normalize the distribution, the inverse logarithm of this reaction time measure is used for analysis such that a more positive measure of the SST corresponds to higher inhibitory control.

#### Visual-motor integration

We assessed visual-motor integration using the Beery VMI 6^th^ Edition (VMI; Beery, 2004), designed for ages 2 to 100 years. The participant’s task is to copy geometric figures. There are specific scoring instructions for each item, resulting in 0 or 1. The test is completed if three consecutive items were failed to be copied correctly. VMI raw scores were used in the analyses as the performance scores for the other cognitive and academic tasks were not adjusted for age.

#### Phonological processing

We assessed phonological processing via the Comprehensive Test of Phonological Processing Second Edition (CTOPP-2; Wagner, Torgesen, Rashotte, & Pearson, 2013). The CTOPP-2 is designed for ages 4:0 through 24:11 and contains a variety of subtests assessing phonological awareness, phonological memory, rapid symbolic naming, and rapid non-symbolic naming. Here we used the Elision subtest, which has been widely used in clinical and typical populations to measure phonological awareness. The participant’s task is to omit a phonological segment (syllable/phoneme) of a word that they previously heard and say the word out loud. The result is another existing word, e.g. “say ‘always’ without ‘all’” [ways] or “say ‘silk’ without /l/” [sick]. Phonological awareness is measured as the sum of all correctly answered items out of a maximum of 34.

### Statistical Analysis

Statistical Analyses were performed with IBM SPSS Statistics (v. 26). Effect sizes were assessed for group comparisons: (partial) η^2^ for non-parametric tests, ANOVAs and ANCOVAs, and Cramer’s V for χ^2^ tests. Because mothers of preterm children reported less education on average then mothers of full-term children, we used maternal education as a covariate in our analyses.

We assessed the mediating effect of cognitive functions on mathematics skills with multiple mediation analyses. A multiple mediation model analyzes if and to what extent the effect of preterm birth on mathematics skills can be accounted for by the effect of preterm birth on cognitive functions, which in turn influence mathematics skills. Direct and mediated effects were evaluated based on their effect size.

The multiple mediation analyses assessed:

1. the total effect model, which considers the total effect of preterm birth on the outcome variable. This does not include the mediators.
2. the direct (unmediated) effect of preterm birth on the outcome variable, when mediators are included.
3. the indirect (mediating) effects of preterm birth on the outcome variable through the mediators.

We used separate multiple mediation models for the three testing times (age 5, 6, and 7 years) and both mathematics skills outcome variables (number and arithmetic skills), resulting in a total of six mediation models. Given the constraints of our relatively small sample, this cross-sectional approach allowed us to remain descriptively close to our data while highlighting differences across the three ages in identical samples.

Due to the intercorrelations of group, maternal education, and cognitive functions, the effect of group can switch from being negative (lower performance in the preterm group) to positive (higher group average in the preterm group, when controlling for other variables in the model). This is because the other variables assume some of the variance that group accounted for when alone in the model.

## Results

### Age and Group differences in mathematics skills and cognitive function

Of primary interest are number and arithmetic skills—our outcome variables. The other two mathematic skill variables—TEMA-3 overall, scaled and percentage—are included for comparison. In addition, the cognitive function variables were analyzed subsequently as potential mediating variables. Group means at the three ages, adjusted for maternal education, are shown for mathematics skills variables in Figure 1 and for cognitive function variables in Figure 2. Table 3 provides statistics for repeated measures, trend analyses of covariance with age as the repeated measure, term status (preterm vs. full-term) as the between-groups factor, and maternal education as the covariate.

**Table 3.** Analysis of Covariance Results for Mathematic Ability and Cognitive Function Scores.

| Variable | Group (PT/FT) |  | Age |  | Age × Group |  |
| --- | --- | --- | --- | --- | --- | --- |
| | $\eta^2$ | *p* | $\eta^2$ | *p* | $\eta^2$ | *p* |
| Mathematics scores |  |  |  |  |  |  |
| TEMA-3 overall (scaled) | .076 | .015 | .031 | .39 | .042 | .074 |
| TEMA-3 overall (%) | .113 | .003 | .429 | <.001 | .005 | .543 |
| Number skills (%) | .104 | .004 | .425 | <.001 | .070 | .020 |
| Arithmetic skills (%) | .145 | .001 | .194 | <.001 | .180 | <.001 |
| Cognitive function scores |  |  |  |  |  |  |
| SWM (inverse) | .030 | .135 | .172 | <.001 | .004 | .609 |
| SST (inverse log) | .096 | .006 | .126 | .002 | <.001 | .940 |
| VMI | .229 | <.001 | .214 | <.001 | .001 | .821 |
| Elision | .139 | .001 | .378 | <.001 | .001 | .886 |

Note. Group, $n = 78$ (51 preterm, 27 full-term). Results are for repeated measures ANCOVA with age (5, 6, and 7 years) as the repeated measure, group as the between-groups factor (PT vs. FT), and maternal education as the covariate. PT: preterm, FT: full-term, TEMA-3: Test for Early Mathematics Ability, 3rd edition, SWM: Spatial Working Memory; SST: Stop Signal Task; VMI: Visual-motor Integration.

**Figure 1.**
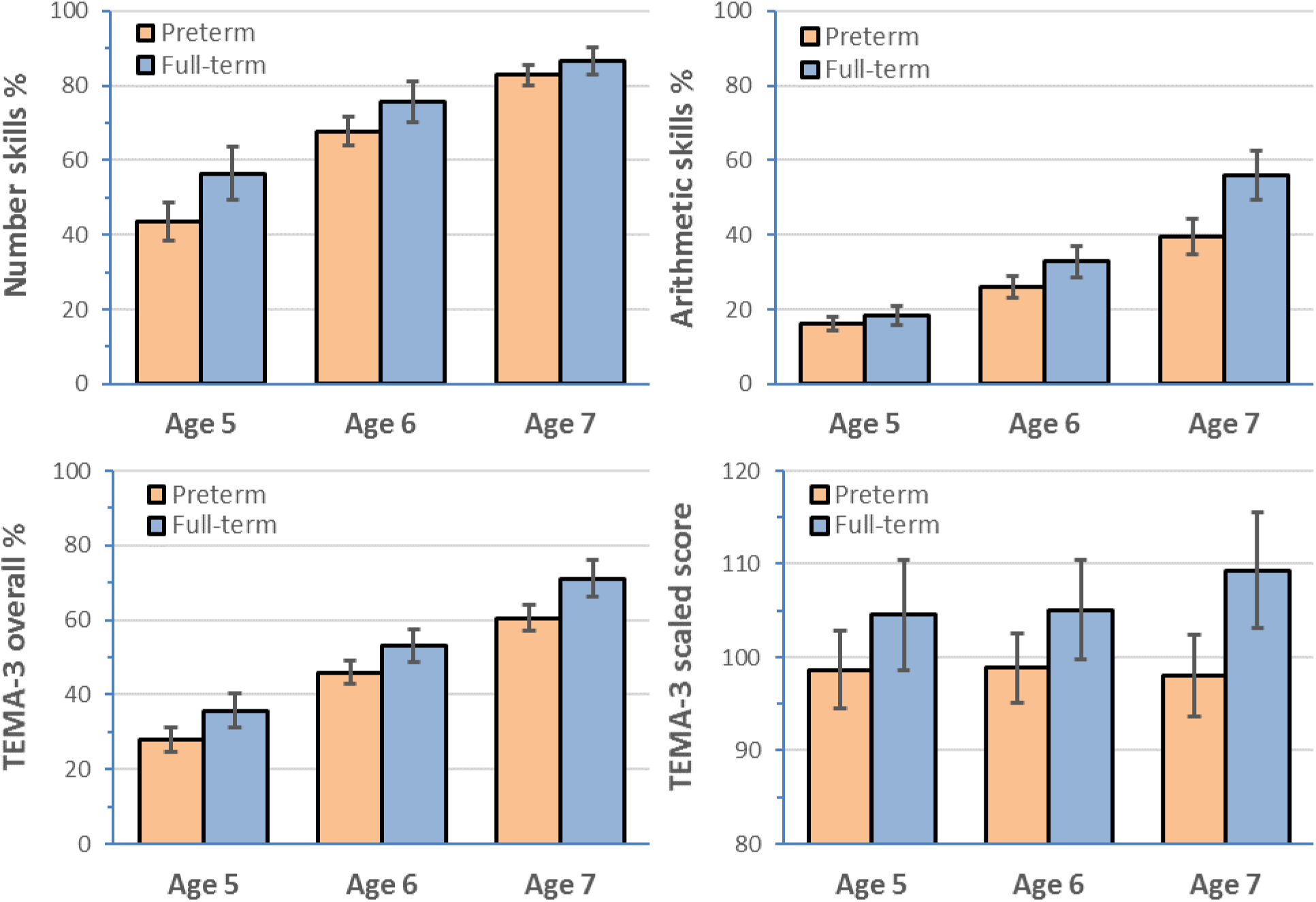
Development of mathematics skills from age 5 to 7 for preterm and full-term groups. TEMA-3: Test for Early Mathematics Ability, 3^rd^ edition. Error bars are 95% confidence intervals.

**Figure 2.**
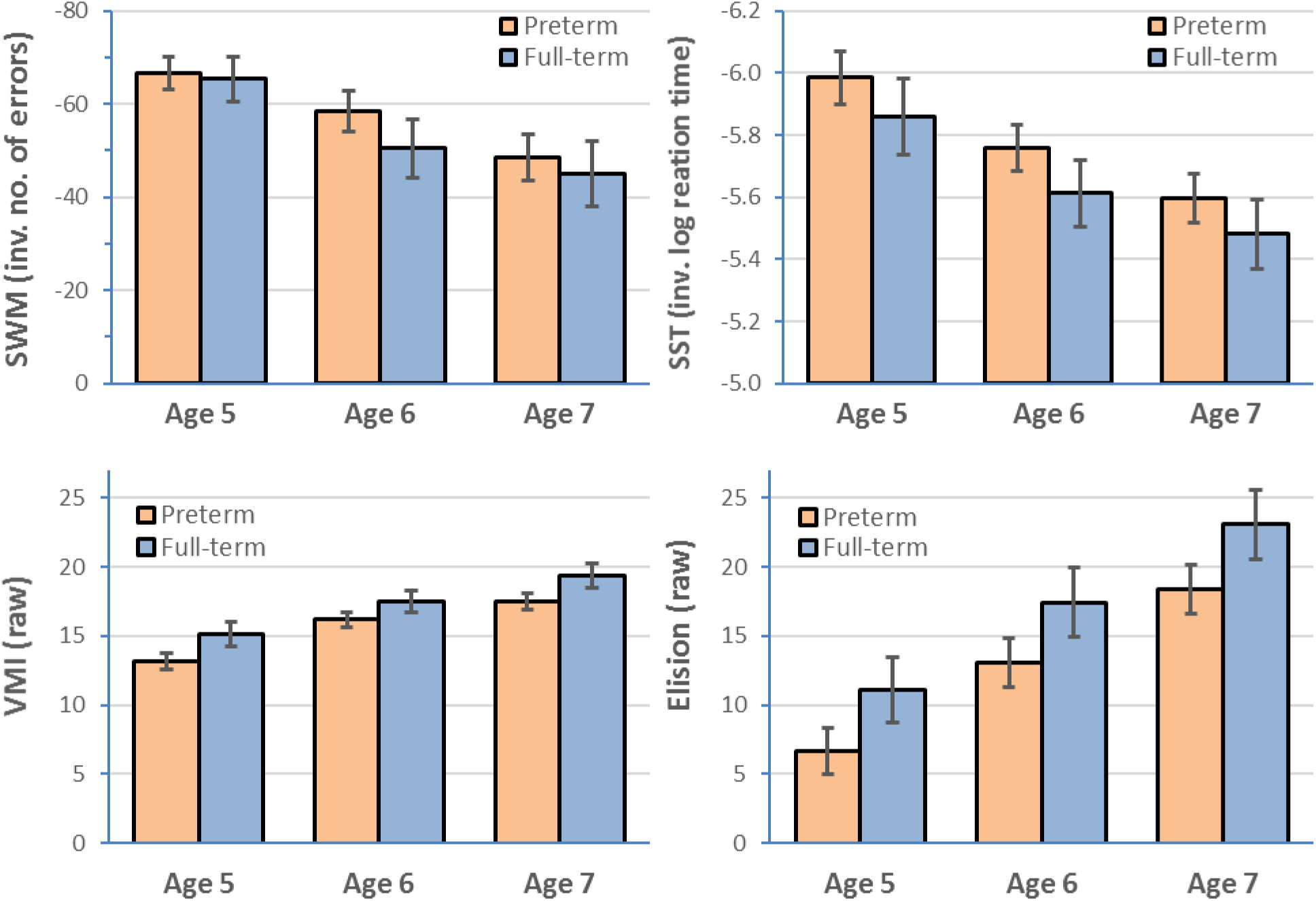
Development of cognitive functions from age 5 to 7 for preterm and full-term groups. SWM: spatial working memory, SST: stop signal task/test of inhibitory control, VMI: visual-motor integration, Elision: test of phonological processing. Error bars are 95% confidence intervals.

Age and group interacted for the number and arithmetic skills: mean differences between preterm and full-term groups decreased with age for number skills but increased with age for arithmetic skills. A repeated measures ANCOVA with type of mathematics skill as additional within-subject-variable found a strong and statistically significant interaction between time, group, and type of skill (F[2,150] = 18.10, *p <* .001, partial η^2^ = .194) —indicating that the difference between the patterns for number and arithmetic skills—decreasing pre-term–full-term differences for number skills but increasing differences for arithmetic skills—was consequential. (The threshold for what we term small or weak, medium or moderate, and large or strong η^2^ is .01, .06, and .14; Cohen, 1988.).

The number skills, arithmetic skills, and TEMA-3 overall raw score increased notably with age with linear trend effect sizes ranging from .19 to .43, *p*s < .001. As expected of a scaled score, the TEMA-3 overall scaled score did not show a linear trend. The mean scores of the preterm children were lower than the means from full-term children for all mathematics skills variables with effect sizes (η^2^) ranging from .076 to .145 (see Figure 1 and, for effect sizes, Table 3).

Scores for cognitive functions increased with age with linear trend effect sizes ranging from .54 to .82, *p*s < .001. The mean scores for preterm children were lower than scores for the full-term children; in particular, the largest effects were observed for VMI, less so for Elision, and SST, and least (and not significantly) for SWM (see Table 3 for effect sizes). Age and group did not interact.

### Non-Parametric Analyses of Number and Arithmetic Skills

Adjusting for maternal education, number skill scores were negatively skewed for the full-term group at age 6 and arithmetic skill scores were positively skewed for the preterm group at all ages (standardized skews > 2.58 absolute). To check whether, apart from extreme scores, number and arithmetic skill scores were reasonably distributed, we examined box-and-whisker plots (see Figure 3). Such plots show distributions graphically, including extreme scores (defined as scores 1.5 times the interquartile range greater than the 75^th^ or less than the 25^th^ percentile; Tukey, 1977). The box-and-whisker plots confirm the pattern shown in Figure 1 that the difference between preterm and full-term medians, which are not influenced by extreme scores, decreased with age for the number skills scores (from 16.7, to 13.4, to 3.6) but increased with age for the arithmetic skills scores (from 0.7, to 6.9, to 19.4) for ages 5, 6, and 7, respectivley. We confirmed these impressions via non-parametric Mann-Whitney U tests showing that *p* values increased from .008, to .014, to .047 for numeric skills but decreased from .052, to < .001, to < .001 for arithmetic skills.

**Figure 3.**
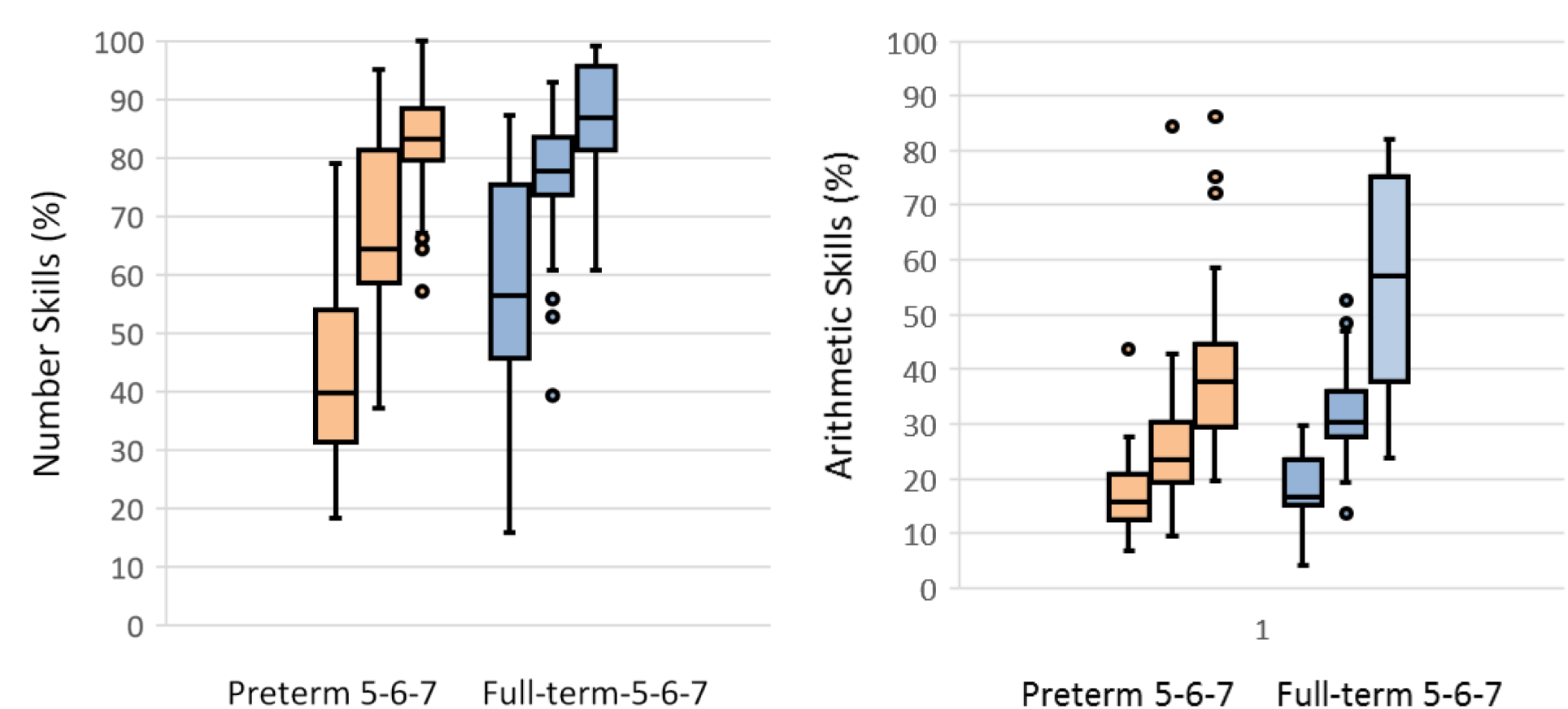
Box-and-whisker plots for preterm and full-term groups from age 5 to 7. The box includes scores from the 25^th^ to the 75^th^ percentile. The whiskers indicate the lowest and highest scores that are not extreme. Extreme scores, defined as any 1.5 times the interquartile range below the 25^th^ or above the 75^th^ percentile, are indicated with circles.

### Simple and Mediation Models Predicting Number and Arithmetic Skills

Models predicting number skills for ages 5, 6, and 7 are shown in Figure 4; similar models predicting arithmetic skills are shown in Figure 5. Displayed first are two single-predictor models, one with term status as the predictor and the other with maternal education as the predictor, thus their path coefficients are simple Pearson correlations coefficients (*r*s). Full-term status was coded 0 and preterm status 1, thus negative coefficients for group signal lower average outcome means for the preterm than the full-term group. The figures also show the percentage of variance not accounted for by each model (i.e., the error variance).

**Figure 4.**
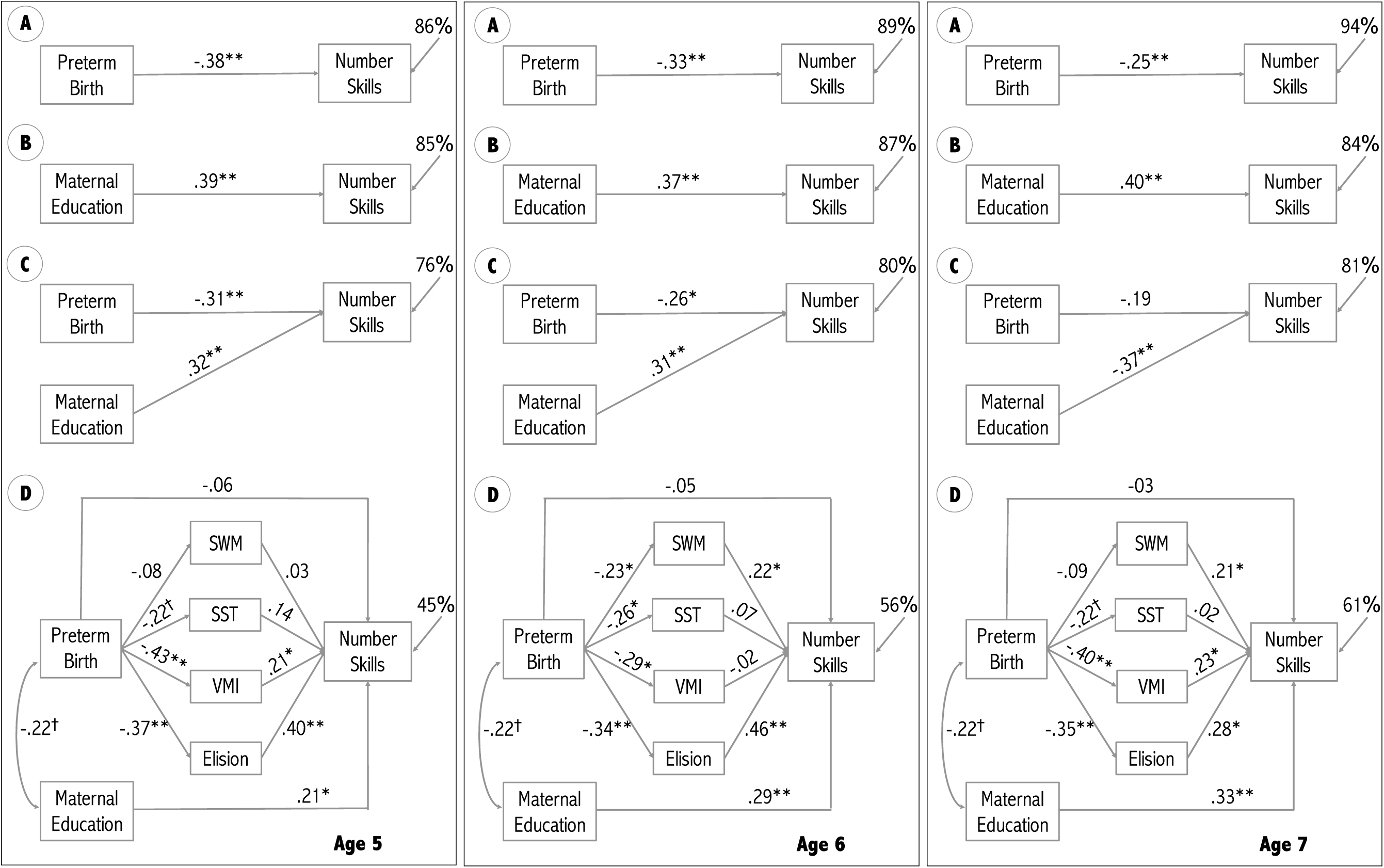
Path diagrams showing the link between group and maternal education, and their effect on number skills over time. A: Effect of preterm birth, B: Effect of maternal education, C: Joint effect of preterm birth and maternal education, D: Mediation models showing the mediating effect of SWM (spatial working memory), SST (stop signal task/inhibitory control), and VMI (visual-motor integration), and Elision (phonological processing) between group and mathematics skills. Coefficients on lines with a one directional arrow represent standardized betas, coefficients on lines with bi-directional arrows are the simple correlation coefficients (*r*). †† *p* < .10, * *p* < .05, ** *p* < .01

**Figure 5.**
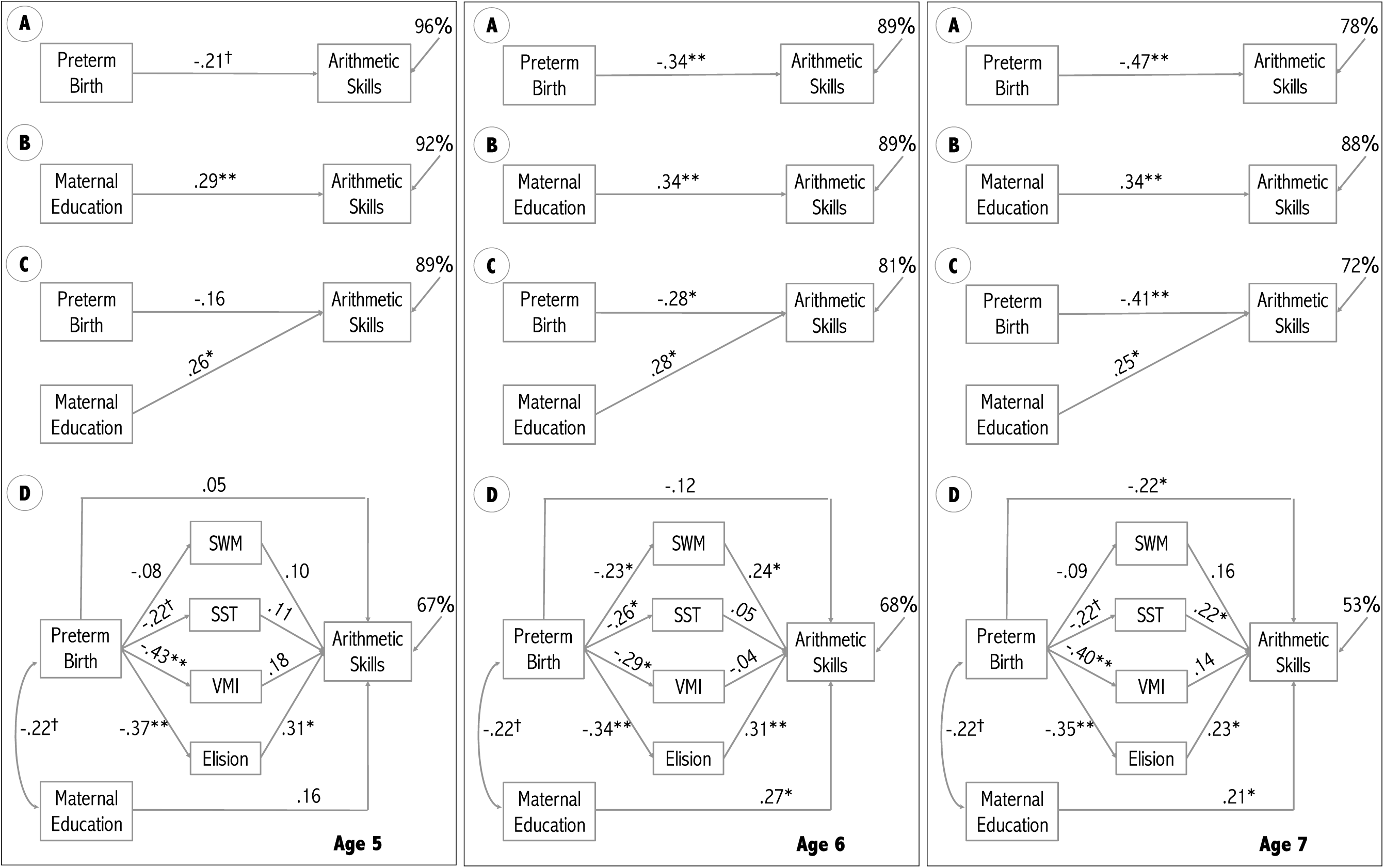
Path diagrams showing the link between group and maternal education, and their effect on arithmetic skills over time. A: Effect of preterm birth, B: Effect of maternal education, C: Joint effect of preterm birth and maternal education, D: Mediation models showing the mediating effect of SWM (spatial working memory), SST (stop signal task/inhibitory control), and VMI (visual-motor integration), and Elision (phonological processing) between group and mathematics skills. Coefficients on lines with a one directional arrow represent standardized betas, coefficients on lines with bi-directional arrows are the simple correlation coefficients (*r*). † *p* < .10, * *p* < .05, ** *p* < .01

Two-predictor, unmediated models—group and maternal education are the predictors— are displayed next; their path coefficients are the partial standardized coefficients of multiple regression (βs). All two predictor models showed redundancy, that is the βs are somewhat smaller than the corresponding *r*s due to the shared and overlapping influence of the two predictors that correlated *r =* .22, *p* = .058, acting in concert. Reflecting the interaction between group, age, and type of mathematics skill noted earlier, group was a stronger predictor of number skills at earlier ages but of arithmetic skills at later ages: the path coefficients decreased with age in magnitude from –.31 to –.26 to –.17 for number skills but increased from –.16 to –.28 to –.41 for arithmetic skills.

Figures 4 and 5 display the mediation models using cognitive functions as the mediators with maternal education as a covariate for number skills and arithmetic skills, respectively. Once mediators were added to the model, the path coefficients for term status declined, becoming inconsequential (< .10 absolute) for number skills and inconsequential, barely small, or small for arithmetic skills at ages 5, 6, and 7, respectively (defining small as .1 to .3 absolute; Cohen, 1988). With age, number skills models accounted for less variance, decreasing from 55% to 44% to 39%, whereas arithmetic skills models accounted for more, from 33% to 32% to 47%, for ages 5, 6, and 7, respectively. In sum, the addition of cognitive functions variables to the models rendered the influence of term status inconsequential, indicating that these cognitive functions variables together mediated most of the influence of term status on number and arithmetic skills.

Table 4 provides the statistics for the mediation models. The table shows how the total effect of term status (i.e., its simple correlation with outcome) can be decomposed into the direct effect of term status (i.e., its β with cognitive function scores and maternal education in the model); the direct, mediated effects of the cognitive functions (i.e., the product of their correlation [*r*] with term status and their partial standardized regression coefficient [β] when term status, other cognitive function scores, and maternal education are included the model); and the direct effect of maternal education (i.e., the product of its *r* with term status and its β when term status and cognitive function scores are included in the model). These direct effects sum to the simple correlation between term status and outcome.

**Table 4.** Mediation Analysis Results for Cognitive Function Scores.

|  | Age 5 |  |  | Age 6 |  |  | Age 7 |  |  |
| --- | --- | --- | --- | --- | --- | --- | --- | --- | --- |
| | r with PT/FT | std. $\beta$ for skill | effect coefficient | r with PT/FT | std. $\beta$ for skill | effect coefficient | r with PT/FT | std. $\beta$ for skill | effect coefficient |
| Number Skills |  |  |  |  |  |  |  |  |  |
| Group |  |  | -.061 (16%) |  |  | -.048 (15%) |  |  | .032 (-13%) |
| SWM | -.08 | .03 | -.002 (.5%) | **-.23\*** | **.22\*** | **-.051 (16%)** | -.09 | .21* | -.019 (7.5%) |
| SST | -.22† | .14 | -.031 (8.2%) | -.26* | .07 | -.018 (5.5%) | -.22† | .02 | -.004 (1.6%) |
| VMI | **-.43\*\*** | **.21\*** | **-.090 (24%)** | -.29* | -.02 | .006 (-1.8%) | **-.40\*\*** | **.23\*** | **-.092 (37%)** |
| Elision | **-.37\*\*** | **.40\*\*** | **.148 (39%)** | **-.34\*\*** | **.46\*\*** | **-.156 (48%)** | **-.35\*\*** | **.28\*** | **-.098 (39%)** |
| M.Ed. | -.22† | .21* | **.046 (12%)** | -.22† | **.29\*\*** | **-.064 (20%)** | -.22† | **.33\*\*** | **-.073 (29%)** |
| Total |  |  | -.378 |  |  | .327 |  |  | .252 |
| Arithmetic skills |  |  |  |  |  |  |  |  |  |
| Group |  |  | .050 (-24%) |  |  | -.119 (35%) |  |  | -.219 (47%) |
| SWM | -.08 | .10 | -.008 (3.8%) | **-.23\*** | **.24\*** | **-.055 (16%)** | -.09 | .16 | -.014 (3.0%) |
| SST | -.22† | .11 | -.024 (11%) | -.26* | .05 | -.013 (3.8%) | **-.22†** | **.22\*** | **-.048 (10%)** |
| VMI | **-.43\*\*** | **.18** | **-.077 (36%)** | -.29* | -.04 | .012 (-3.6%) | **-.40\*\*** | **.14** | **-.056 (12%)** |
| Elision | **-.37\*\*** | **.31\*** | **-.115 (55%)** | **-.34\*\*** | **.31\*\*** | **-.105 (31%)** | **-.35\*\*** | **.23\*** | **-.081 (17%)** |
| M.Ed. | -.22† | .16 | -.035 (17%) | -.22† | **.27\*** | **-.059 (18%)** | -.22† | **.21\*** | **-.046 (10%)** |
| Total |  |  | -.211 |  |  | -.337 |  |  | -.466 |

Note. Group, n = 78 (51 preterm, 27 full-term). Mediating effects of .04 or greater are bolded. M.Ed: maternal education
† p < .10, \* p < .05, \*\* p < .01

We are now able to ask which particular mediators are noteworthy. One criterion is that both their constituent *r* and β coefficients need to be statistically significant (Baron & Kenny, 1986). Ten of the 24 mediating effects of the cognitive functions shown Table 4 meet the *p* < .05 for both *r* and β criterion. However, this criterion, like statistical significance, depends on sample size. A criterion based on absolute effect size might be a better choice (Wilkinson, 1999). We suggest viewing mediating effects of .04 or larger as worthy of further consideration. Three additional mediating effects have an effect size larger than .04: One mediating effect with *p* < .10 for the *r*, and < .05 for the β, and two mediating effects with *p* < .01 for the *r* but was .132, and .155 for the β. These 13 mediating effects of cognitive functions are bolded in Table 4.

The mediator with the largest and most consistent direct effects was Elision; its direct effects averaged .117, were larger for number skills, and decreased with age. For number skills, the absolute effect coefficients accounted for 39%, 48%, and 39% of the total effect at ages 5, 6, and 7, respectively. VMI accounted for 24% of the total effect at age 5, and 37% at age 7, while the mediating effect was marginal at age 6. The mediating effects of SWM and SST on number skills were small at all time points, though at age 6 SWM had a significant mediating effect of –.051 (16%). The effect of maternal education on number skills increased over time both in absolute and relative size. Maternal education contributed 12%, 20%, and 29% of the total effect, respectively for ages 5, 6, and 7.

In contrast to number skills, the total and direct effect of group on arithmetic skills increased over time. Strikingly, the direct effect of group on arithmetic skills at age 7 accounted for 47% of the total effect. Like number skills, phonological processing is the strongest mediator among the cognitive functions included in the model. The mediating effect of phonological processing is largest in absolute and relative size at age 5 and decreases over time (their absolute effect coefficients account for 55%, 31%, and 17% of the total effect at ages 5, 6, and 7, respectively). Following the same pattern as for number skills, VMI and SST showed mediating effects at age 5 and 7, and SWM at age 6. The effect of maternal education on arithmetic skills accounted for 10–18% of the total effect of preterm birth.

### Sex Effects

Sex was not included as a variable in the mediation model as the number of boys and girls did not significantly differ between the groups. Nonetheless, to explore sex effects, we reanalyzed the previous trend analyses of number and arithmetic skills including sex as a between-subjects variable. The pattern of male–female differences differed by birth status; group by sex interaction *F*(1,74) = 4.72 and 4.24, 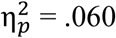 and .054, *p* = .022 and .043, for number and arithmetic skills, respectively. Averaging across the three ages, preterm boys scored lower than girls on number skills (63% vs. 67%) and about the same on arithmetic skills (27% vs. 28%). In contrast, full-term boys scored higher than girls on both number (76% vs. 69%) and arithmetic skills (46% vs. 421%).

We considered whether the group by age interaction described earlier might have been moderated by sex. For number skills the answer is no. The strength of the age by group interaction was moderate and statistically significant, the same as for the numeric skills analysis shown in Table 3; 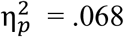 and .070, *p* = .023 and .020, respectively. But both the age by sex and the age by group by sex interactions were small or less in magnitude and not statistically significant; 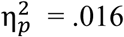 and ∼0, *p* = .27 and .96, respectively.

The answer is more complex for arithmetic skills. The age by group interaction was statistically significant, the same as for the arithmetic skills analysis shown in Table 3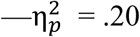 and .18, *p* = < .001 for both, respectively. For arithmetic skills, however, both the age by sex and the age by group by sex interactions were moderate and statistically significant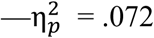 and .11, *p* = .019 and .004, respectively. Follow-up analyses for each sex separately showed that the group by age interaction was statistically significant for boys but not statistically significant in girls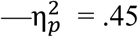 and .015, *p* = < .001 and = .46, respectively. The mean for boys in arithmetic skills showed divergence, increasing from 16% to 27% to 39% for preterm participants and from 18% to 36% to 65% for full-term participants. Means for girls did not diverge, increasing from 17% to 25% to 40% for preterm and from 18% to 29% to 45% for full-term children.

## Discussion

Preterm birth can have a significant and negative impact on mathematics achievement. Our first aim was to shed light on the impact of preterm birth on two distinct kinds of mathematics skills: number and arithmetic skills. Analyses according to this division showed that there were considerable differences in the developmental trajectories both in the development of these skills *per se*, and in the development of these skills in preterm and full-term children. Only number and arithmetic skills showed a significant interaction effect of term status and time; this was neither reflected in overall mathematics ability nor in any of the assessed cognitive functions.

The poorer performance of preterm children compared with full-term children in number skills was largest at age 5 and decreased over time. Following preterm birth, children need longer to reach the same level of proficiency in number skills compared to children who were born full-term. That is, the deficit in counting and comparing magnitudes of numbers is not persistent. Instead preterm children are delayed in their development of such skills but eventually “catch up.” A qualitatively similar developmental trajectory with a “catch-up” in performance across time has been reported for executive functions and receptive vocabulary in children born preterm (Luu et al., 2009; Ritter, Nelle, Perrig, Steinlin, & Everts, 2013). To our knowledge, this is the first study of preterm children showing a “catch-up” in number skills performance. Some studies suggest that number skills continue to develop into adolescence (Lyons et al., 2014). Note that the dependent variable in those studies were reaction times rather than accuracy, and the TEMA-3 does not measure reaction times. Thus, our findings for a maturing of accuracy measures at age 7 does not contradict potential continued refinement of number skills measured in reaction time. Together, these studies hint towards a maturational delay in various skills following preterm birth. The underlying reason may be maturational delay of brain structures, specifically white matter tracts vulnerable to damage in very preterm birth (Volpe, 2009). In support, MRI studies of preterm children and full-term controls found that brain development in both groups follows a similar trajectory that is delayed for children who were born preterm (Sripada et al., 2018).

In contrast, there was an increase in the performance gap in arithmetic skills with age. Lower arithmetic skills may arise from lower number skills if number skills act as a scaffold to promote learning of arithmetic skills. Perhaps children born very preterm that show a delay in the typical acquisition of number may eventually “catch up” in their arithmetic skills. This seems unlikely. Others have found poorer than expected mathematics achievement among older children and adolescents who were born preterm (Akshoomoff et al., 2017; Litt et al., 2012; Taylor et al., 2009). Similar to our findings, a meta-analysis of 17 studies found that arithmetic performance of preterm children and adolescents (6–18 years of age) was 0.71 SD below full-term controls (Twilhaar, Kieviet, Aarnoudse-Moens, Elburg, & Oosterlaan, 2018). This hints towards persistent lower performance in complex mathematics skills following preterm birth. A persistent effect of preterm birth on arithmetic skills may be explained by persistent deficits in cognitive functions that contribute to these skills.

Our second aim sought to identify the specific cognitive functions that mediate number and arithmetic skills as children move from preschool through first grade. We used a mediation model, testing for the potentially mediating effect of spatial working memory, inhibitory control, visual-motor integration, and phonological processing. The assessment of the variety of skills, and analysis via multiple mediation models has several advantages: indirect effects of variables can be compared in the context of others, the direct effect of group can be determined, and the parameter estimation is more accurate. Importantly, theoretical causality drives mediation models, but does not provide proof of causality. The direction of influence here was determined by using non-academic cognitive functions (spatial working memory, inhibitory control, phonological processing, visual-motor integration) as mediators of the mathematics skills. The structure of our mediation models are in line with our previous study of 5-year old children born preterm (Hasler & Akshoomoff, 2019) and studies of children with spina bifida (Barnes et al., 2014; Raghubar et al., 2015).

Phonological processing was the strongest mediator for both number skills and arithmetic skills. This is consistent with previous studies showing the effect of early phonological skills on mathematics performance. Phonological awareness in preschool and kindergarten has been reported to be predictive of several distinct mathematics skills, including numeration and calculation (LeFevre et al., 2010). Phonological awareness has also been associated with arithmetic skills in older children (De Smedt et al., 2010), and shown to mediate the relationship between other developmental disorders (spina bifida) and calculation skills (Barnes et al., 2014). This strong association may be explained by a shared network of brain regions associated with both phonological processing and mathematics skills. The temporo-parietal cortex, specifically the arcuate fasciculus are candidate regions. For example, Van Beek, Ghesquière, Lagae, & De Smedt (2014) found a correlation between the arcuate fasciculus microstructure and children’s addition/multiplication skills. These effects disappear when covarying for phonological processing, pointing towards an involvement of phonological processing when solving mathematics problems.

In line with previous research, we found that visual-motor integration measured by the Beery VMI, was a strong mediator for both types of mathematics skills at age 5 and age 7. Performance on the VMI has been associated with mathematics skills in children at age 4 (Verdine, Irwin, Golinkoff, & Hirsh-Pasek, 2014), and 5-18 years (Carlson, Rowe, & Curby, 2013). This link has rarely been investigated in children born preterm. Perez-Roche et al. (2016) found an effect of visual abilities on school performance in small for gestational age children (though school performance was assessed with a parent questionnaire). Others have found a link between low motor and visuospatial function and low academic achievement in extremely preterm children (Marlow, Hennessy, Bracewell, & Wolke, 2007).

Inhibitory control had a mediating effect on arithmetic skills at age 7. In line with these findings, inhibitory control has been reported to be associated with procedural arithmetic skills in children and conceptual knowledge in adults (Gilmore, Keeble, Richardson, & Cragg, 2015). Inhibitory control has also been shown to be predictive of early mathematics skills in 3-5 year old children (Blair & Razza, 2007) and of arithmetic skills in fourth graders (Passolunghi & Siegel, 2001). However, these studies investigated the role of inhibition on mathematics in the absence of other cognitive functions, which is likely to account for different results.

The participants in the full-term group had higher maternal education than the preterm group. This may be due to them being a convenience sample. However, this might also be a closer representation of the true distribution of maternal education in the preterm and full-term population (Behrman & Butler, 2007; Thompson, Irgens, Rasmussen, & Daltveit, 2006). Higher maternal education has been linked to better mathematics and reading performance, fewer behavioral problems, and less grade repetition (Carneiro, Meghir, & Parey, 2013; ElHassan et al., 2018). We included maternal education in the mediation model and, as expected, higher maternal education showed a significant positive effect on both number and arithmetic skills.

One might expect that the influence of social variables, such as maternal education, on mathematics would increase over time, and conversely that the influence of biological variables associated with prematurity would decrease. However, in a study of extremely preterm children 8 and 18 years of age, Doyle et al. (2015) found that the adverse effect of preterm birth persists over time. In line with their results, we found that the effect of group on arithmetic skills increased over time. Interestingly, not only the total effect of preterm birth, but also the direct effect increases from age 5 years to 7 years. In fact, at age 7 years only about half of the total effect of preterm birth on arithmetic skills was mediated by other cognitive functions. In contrast, the direct effect of preterm birth on number skills was statistically non-significant at all three time points, meaning the total effect of preterm birth was fully mediated by cognitive functions (mainly phonological processing and visual-motor integration).

In addition, at age 7, we found an interaction effect of sex and group, with males having significantly higher arithmetic skills scores than females in the full-term group only. There is a vast body of literature on gender differences and learning and education. Studies have shown that teachers’ implicit gender bias affects achievement in children, and that children themselves develop gender stereotypes of males being “smarter” and “better in math” already in elementary school (Bian, Leslie, & Cimpian, 2017; Cvencek, Meltzoff, & Greenwald, 2011; Lavy & Sand, 2015). One possible explanation as to why we did not observe this effect in the preterm group might be that the differences in cognitive processing related to preterm birth do not effectively influence preterm males; that is, societal and cultural influences do not impact preterm children in the same way as full-term children. On the other hand, it might be that these influences are delayed in preterm children and may become evident later in their educational development.

A significant takeaway message from this study is that our healthy preterm children showed reliable adverse effects of preterm birth. Although severity of neonatal brain pathology is linked to cognitive outcomes in preterm children (Murray et al., 2014), preterm children on the healthy end of the continuum still show significant cognitive disparities that negatively impact their academic performance. The lack of severe disabilities is not an indication of typical developmental outcomes and any child born preterm requires assessment before formal education begins, with continued monitoring.

## Conclusion

Inclusion of a broad variety of cognitive functions distinguishes this study and provides an important contribution to the literature. Previous work on the effect of cognitive functions on mathematics skills have primarily focused on functions within a single cognitive domain. Cognitive functions and academic skills in children develop dynamically between ages 5 and 7 and change in complexity. Our study gives valuable insight into how these skills are impacted by preterm birth and the contribution of cognitive functions on mathematics skills in both full-term and preterm birth children. Neuropsychological assessment of preterm children at 5 years of age has been shown to be predictive of cognitive functions and need for educational support at age 11 (Lind, Nyman, Lehtonen, & Haataja, 2019). Hence, the cognitive functions investigated here, particularly phonological processing, can be neuropsychological markers for early evaluation of children with increased risk for difficulties in mathematics.

## Acknowledgments

The Eunice Kennedy Shriver National Institute of Child Health and Human Development grant R01HD075765 supported this work. We thank the children and families who participated in this study. Thank you to Holly Hasler, Ph.D., Stephanie Torres, M.A., Kelly McPherson, Akshita Taneja, and Rubaina Dang; and our collaborators: Martha Fuller, Ph.D., PPCNP-BC, Yvonne Vaucher, M.D., Terry Jernigan, Ph.D., and Joan Stiles, Ph.D.

## Disclosure of Interest

The authors report no conflict of interest.

## Data Availability Statement

The data supporting the findings of this study are available from Dr. Natacha Akshoomoff upon reasonable request.

## Notes

### Competing Interest Statement

The authors have declared no competing interest.

### Summary of Updates

Expansion of the results section, including one additional figure Clarification of introduction and methods

